# Pixel-Precise Lesion Localization in WSIs via Weakly Supervised Streaming Convolution with ReLSE and Adaptive Self-Training

**DOI:** 10.1101/2024.10.03.616445

**Authors:** Chi-Chung Chen, Yi-Chen Yeh, Matthew MY Lin, Chao-Yuan Yeh

## Abstract

A robust artificial intelligence-assisted workflow for tumor assessment in pathology requires not only accurate classification but also precise lesion localization. While current weakly supervised learning methods significantly reduce the need for extensive annotations and leverage large quantities of annotation-free whole-slide images (WSIs) to enhance classification robustness, they often fall short in segmentation accuracy. We attribute this limitation to the optimization goals in classification, which tend to focus solely on the most representative features—an approach that is particularly inefficient for WSIs with gigapixel resolution. To address this challenge, we introduce a novel approach based on streaming convolution, an end-to-end method for WSI training. Our contributions include the Rectified LogSumExp (ReLSE) pooling method and adaptive pseudo annotation generation for self-training, both designed to encourage models to learn from sub-representative features. Using only slide-level annotations from the CAMELYON16 dataset, our method achieves a significant improvement in metastasis localization, with a patch-level recall from 53.88% to 79.18% at a precision of 80%. This conclusion also holds for a dataset collected from Taipei Veterans General Hospital (TVGH), used in the assessment of lung cancer lymph node metastasis with a recall improved from 42.98% to 72.27%. The proposed method can also be integrated with additional manual annotations. Experiment results showed the semi-supervised model (recall: 91.12%) could even outperform the strongly supervised model (recall: 81.80%) on the TVGH dataset.

## Introduction

The application of deep learning in digital pathology has gained traction in clinical settings, as numerous studies have demonstrated that AI-assisted workflows can enhance diagnostic accuracy and reduce human workload.^1–3^ AI-assisted tumor assessment workflows, such as those used for metastasis localization in gastric lymph nodes, have been shown to improve diagnostic accuracy and decrease pathologists’ review time.^1^ These workflows require both high classification accuracy and precise lesion localization. However, training lesion localization models typically demand detailed annotations, which are labor-intensive and restrict dataset size. The transition from models that fit well with collected datasets to robust, real-world medical devices presents a significant challenge, particularly with regard to model generalization across diverse patient cohorts. The need for large, diverse datasets is fundamental to expanding the generalization capacity of models. Unfortunately, the high cost and complexity of acquiring detailed annotations limit the development of strongly supervised learning approaches. As a result, advancing weakly supervised, which requires no detailed annotations, has become increasingly important.

Weakly supervised learning for whole-slide images (WSIs) presents unique challenges compared to natural image applications, primarily due to the gigapixel resolution of WSIs. The dominant paradigm for WSI analysis is multiple-instance learning (MIL)^4^, which divides a WSI into patches to maintain a manageable memory footprint. MIL generally consists of two stages: intra-patch feature extraction and inter-patch aggregation. While feature extraction can be enhanced using self-supervised pretraining^5–7^, aggregation methods have evolved from simple maximum or mean pooling to more sophisticated approaches. Techniques such as recurrent neural networks (MIL-RNN)^8^ and attention mechanisms (e.g., CLAM^9^, ABMIL^10^, DSMIL^11^, TransMIL^12^) have been introduced to improve inter-patch aggregation. A key limitation of most MIL approaches is the separate training of feature extraction and aggregation models, often with frozen feature extraction weights, thus hindering the refinement of feature extraction based on slide-level labels. In contrast, streaming convolution^1,13–15^ utilizes gradient checkpointing^16^ and online patching, enabling end-to-end convolutional neural network (CNN) training on gigapixel images. This allows both feature extraction and non-local information aggregation to be jointly optimized through backpropagation, which explains its superior performance when applied to large datasets. Similar to MIL, the aggregation of spatial information plays a crucial role in streaming convolution. Although global average pooling (GAP) is a common choice for the final pooling layer in CNNs, it has been shown to be suboptimal for WSI training.^17^ In contrast, global maximum pooling (GMP), which emphasizes the most significant features in the final feature map, has been particularly effective for detecting small lesions.^1,13,15,17^ Additionally, attention-gated classification heads have been employed to learn optimal aggregation weights.^14^

In this study, we introduced an efficient pooling method for streaming convolutional WSI training, termed Rectified LogSumExp (ReLSE). ReLSE operates between GMP and global sum pooling (GSP), striking a balance between completely ignoring non-significant features and fully incorporating them. By adjusting the positive hyperparameter γ, ReLSE can be tuned to lean toward GMP (when γ is large) or GSP (when γ approaches 0^+^). We evaluated our method in terms of both classification and localization performance against state-of-the-art approaches as well as strongly supervised learning techniques. To assess generalization, we employed several datasets, including breast cancer lymph node metastasis datasets, Camelyon16^18^ and Camelyon17^18^, The Cancer Genome Atlas – Non-Small Cell Lung Cancer (TCGA-NSCLC; see Supplementary for the results), and a 3,024-LN lung cancer lymph node dataset collected from Taipei Veterans General Hospital (TVGH).

## Results

### Classification of breast cancer metastasis in sentinel lymph nodes

We first evaluated our method using the CAMELYON16 dataset, comparing its performance against other weakly supervised approaches. For a fair comparison, we used only slide-level labels (metastatic or negative), disregarding the detailed annotations provided by the dataset. Trained at a resolution of 0.5 μm/pixel, the streaming CNN with a ResNet-50 backbone and global max pooling (GMP) achieved an area under the receiver operating characteristic (ROC) curve (AUC) of 0.9416 (95% CI: 0.8929–0.9904), as shown in Figure 1. With a 0.5 decision threshold, the model reached a sensitivity of 85.42% and a specificity of 91.36%. In comparison, the streaming CNN with ResNet-50 and the proposed ReLSE pooling significantly improved the AUC to 0.9633 (95% CI: 0.9234–1.0000). At the same cut-off of 0.5, the sensitivity remained similar at 85.71%, while the specificity increased to 97.50%. Despite varying experimental settings, our classification performance is competitive with other state-of-the-art methods, including DTFD-MIL (AUC: 0.946; AFS; ResNet-50; 0.5 μm/pixel)^19^, IBMIL (AUC: 0.8951; DTFD-MIL MaxS; ResNet-18; 0.5 μm/pixel)^20^, and StreamingCLAM (AUC: 0.9757; ResNet-34; 0.5 μm/pixel)^14^.

**Figure 1:**
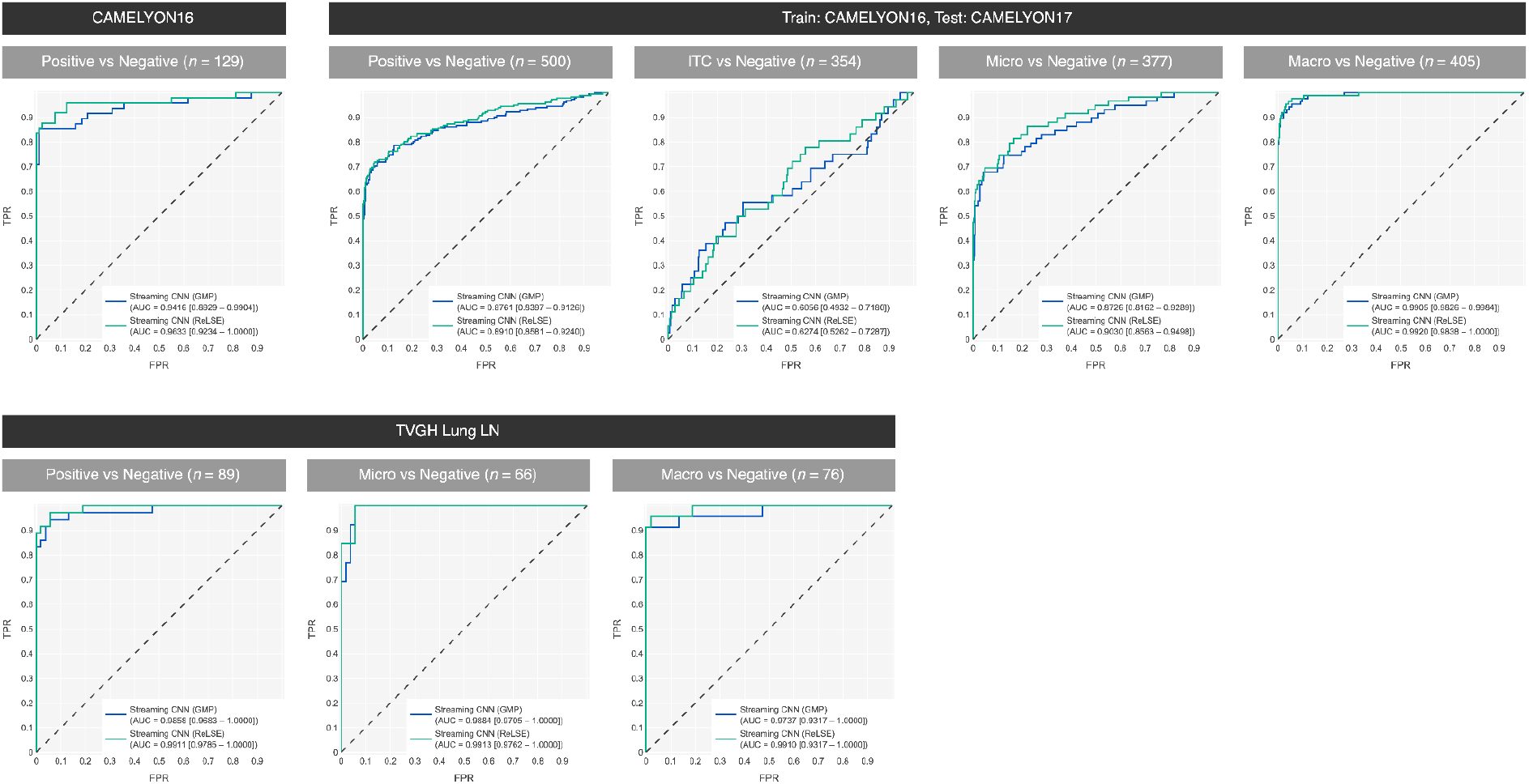
ROC curves of streaming CNN classifiers on different datasets to compare the effectiveness of GMP and ReLSE.

We further examined the effect of lesion size using the CAMELYON17 dataset, where each of the 500 slides is labeled as negative, isolated tumor cells (ITC; lesion size < 0.2 mm), micrometastasis (micro; lesion size 0.2–2 mm), or macrometastasis (macro; lesion size > 2 mm). The overall AUC for differentiating positive from negative slides using the streaming CNN with GMP was 0.8761 (95% CI: 0.8397–0.9126), whereas the streaming CNN with ReLSE achieved a slightly higher AUC of 0.8910 (95% CI: 0.8581–0.9240). By excluding all positive cases except ITC, we found the AUC for distinguishing ITC from negative slides to be 0.6056 (95% CI: 0.4932–0.7180) for the GMP model, and 0.6274 (95% CI: 0.5262–0.7287) for the ReLSE model. Similarly, for micro vs negative cases, the AUC for the GMP model was 0.8726 (95% CI: 0.8162–0.9289), while the ReLSE model improved this to 0.9030 (95% CI: 0.8563–0.9498). In the macro vs negative classification, both models performed similarly, with AUCs of 0.9905 (95% CI: 0.9826–0.9984) and 0.9920 (95% CI: 0.9838–1.0000) for GMP and ReLSE, respectively. These stratified analyses indicate that the ReLSE model’s superiority primarily stems from its enhanced ability to detect smaller lesions, such as ITC and micrometastases.

### Lesion localization of breast cancer metastasis in sentinel lymph nodes

A successful AI-assisted pathology workflow requires not only precise classification but also accurate lesion localization for efficient validation. First, we qualitatively compared different methods, as shown in Figure 2. These methods include visualizing streaming CNNs with GMP and ReLSE pooling using class activation mapping (CAM)^21^, and employing self-training with semantic Feature Pyramid Networks (FPN)^22,23^ based on pseudo-labels generated from CAM results. For each method, a threshold was selected to maintain a patch-level precision of 80% with a patch size of 64 × 64 μm^2^, ensuring a fair comparison without bias toward either high precision or recall. CAM serves as an off-the-shelf visualization tool for streaming CNNs. From qualitative observations, streaming CNNs utilizing ReLSE pooling demonstrated superior performance in identifying smaller lesions compared to those using GMP pooling, particularly when paired with both CAM and FPN. Additionally, the self-trained FPNs significantly enhanced the sensitivity of tumor lesion localization, regardless of lesion size.

**Figure 2:**
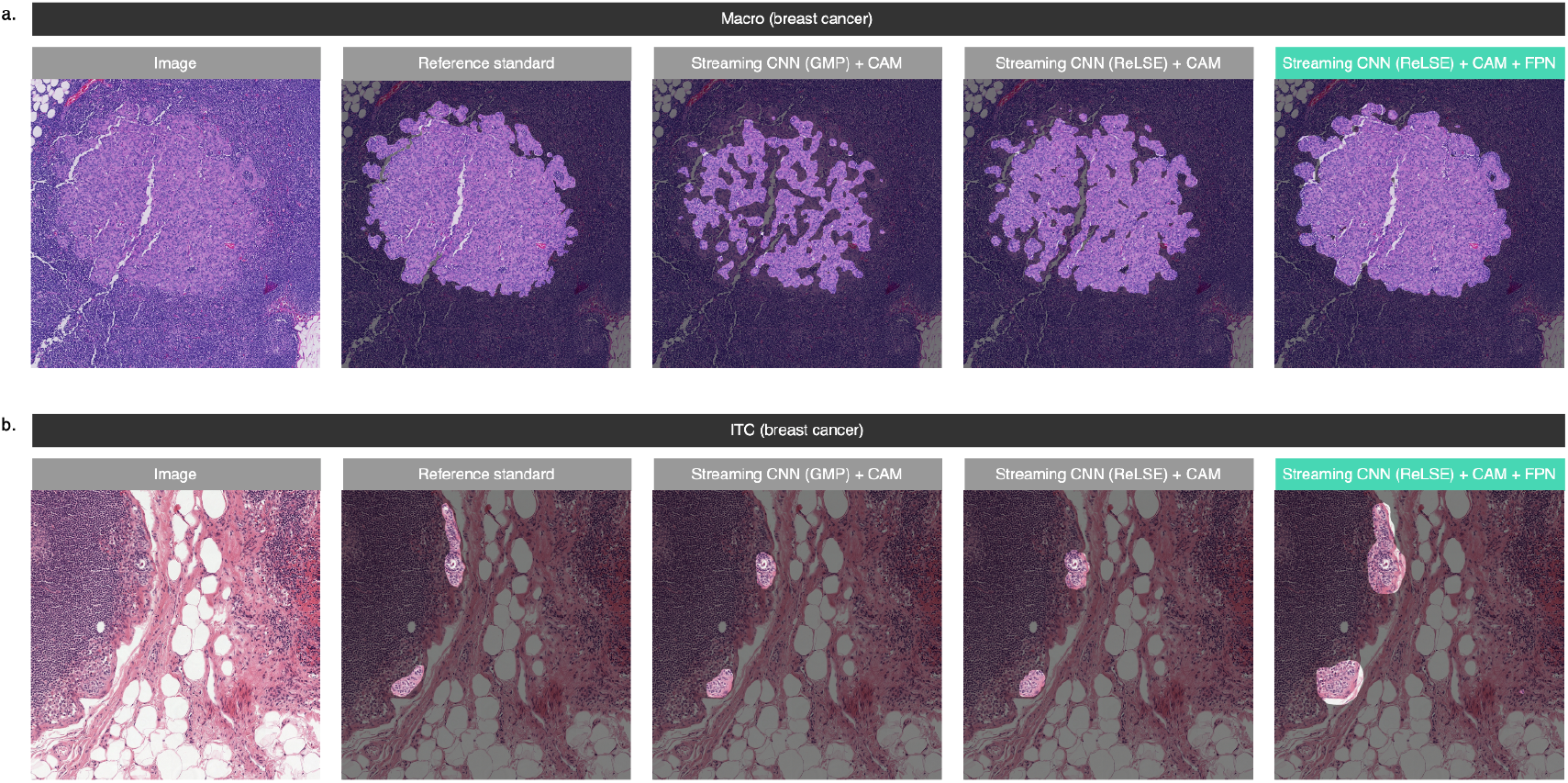
Breast cancer LN metastasis lesion localization of various models on the CAMELYON16 dataset. (a) Image of macro and the identification results of various models. (b) Image of ITC.

Next, we conducted a quantitative evaluation using the 128 test WSIs from the CAMELYON16 dataset, excluding one WSI due to incomplete annotations. Thresholds were again set to achieve a fixed patch-level precision of 80%. Patch-level recalls were computed with a patch size of 64 × 64 μm^2^. To assess the ability of models to detect lesions of varying sizes, annotations were categorized into ITC, micro, and macro. For each category, patch-level recall was defined as the proportion of true positive patches within the annotated areas, while lesion-level recall referred to the proportion of true positive annotations with highlighted regions of any size. When accurate delineation of lesion areas takes precedence over maximizing the number of detected lesions, patch-level recall provides a more representative measure than lesion-level recall, and vice versa. The quantitative results are summarized in Table 1. In summary, both ReLSE pooling and self-training enhanced overall sensitivity and consistently improved lesion localization across the ITC, micro, and macro lesion categories. Notably, the combination of ReLSE pooling and self-training significantly boosted overall recall (79.18% compared to 53.88%) and more than doubled recall for ITC identification (51.85% compared to 21.79%) when compared to the baseline approach using GMP pooling and CAM.

**Table 1:** Performance of Lesion Localization for Breast Cancer Metastasis in Sentinel Lymph Nodes. Four weakly supervised streaming CNN models were trained on 270 Camelyon16 training WSIs using only slide-level labels. Two semi-supervised FPN mixed pseudo labels from ReLSE-CAM-FPN with 20% and 100% of manual annotations, respectively. A strongly supervised FPN was trained using manual annotations. These models were evaluated on 128 test WSIs with exhaustive annotations. Each patch measures 64 × 64 μm^2^. Patch-level recall is defined as the ratio of true positive patches to the total number of positive patches. A lesion-level recall is determined by the presence of any predicted positive patches within a lesion area, classifying it as a predicted positive lesion.

|  |  | Patch-level Recall |  |  |  | Lesion-level Recall |  |  |
| --- | --- | --- | --- | --- | --- | --- | --- | --- |
| Model | Precision | All | ITC | Micro | Macro | ITC | Micro | Macro |
| Streaming CNN (Weakly supervised learning) |  |  |  |  |  |  |  |  |
| GMP-CAM (baseline) | 0.8000 | 0.5388 | 0.2179 | 0.5375 | 0.5438 | 0.3048 | 0.8321 | **1.0000** |
| ReLSE-CAM | 0.8000 | 0.5505 | 0.2434 | 0.5645 | 0.5546 | 0.3426 | 0.8577 | **1.0000** |
| GMP-CAM-FPN | 0.8000 | 0.6341 | 0.3004 | 0.6554 | 0.6383 | 0.3583 | 0.8431 | **1.0000** |
| ReLSE-CAM-FPN (proposed) | 0.8000 | 0.7918 | 0.5185 | 0.8147 | 0.7950 | 0.5678 | 0.9343 | **1.0000** |
| Semi-supervised learning |  |  |  |  |  |  |  |  |
| FPN (20% annotations; proposed) | 0.8000 | 0.8449 | 0.5661 | **0.8156** | 0.8506 | 0.6401 | **0.9416** | **1.0000** |
| FPN (100% annotations; proposed) | 0.8000 | **0.8778** | **0.6431** | 0.8064 | 0.8848 | **0.7223** | 0.9088 | **1.0000** |
| Strongly supervised learning |  |  |  |  |  |  |  |  |
| Strongly supervised FPN | 0.8000 | 0.8772 | 0.6095 | 0.7993 | **0.8849** | 0.7001 | 0.9015 | **1.0000** |

Additionally, we incorporated partial (20%) and full (100%) sets of detailed annotations with pseudo annotations to train models in a semi-supervised manner. For slides containing both detailed and pseudo annotations, we merged their contours to generate the training annotations for the semi-supervised models. The experimental results demonstrated that adding annotations substantially improved recall over the weakly supervised model. When all detailed annotations were utilized, the overall recall was comparable to that of a purely strongly supervised model. However, the inclusion of pseudo annotations led to a slight improvement in ITC detection, increasing from 60.95% to 64.31%.

### Classification of lung cancer metastasis in lymph nodes

We further evaluated our training pipeline on an in-house dataset of lymph node specimens from lung cancer patients, collected at TVGH. The model was trained using 3,024 LN images annotated from 413 WSIs at a pixel spacing of 1.0 μm. On a test set comprising 89 slides, the model utilizing GMP achieved a slide-level AUC of 0.9858 (95% CI: 0.9683–1.0000). In contrast, the model incorporating ReLSE demonstrated superior performance, with an AUC of 0.9911 (95% CI: 0.9785–1.0000).

For a stratified analysis, we further subdivided positive cases into micro and macro. Interestingly, the GMP-based model showed better performance on micro, achieving an AUC of 0.9884 (95% CI: 0.9705–1.0000) when distinguishing micro from negative cases, compared to an AUC of 0.9737 (95% CI: 0.9317–1.0000) for macro vs negative. The ReLSE-based model, however, exhibited consistent performance across both subgroups, achieving AUCs of 0.9913 (95% CI: 0.9762–1.0000) for micro and 0.9910 (95% CI: 0.9740–1.0000) for macro. These results demonstrate ReLSE’s ability to enhance the detection of both small and large lesions, outperforming the GMP model in both cases.

### Lesion localization of lung cancer metastasis in lymph nodes

The lesion localization performance was evaluated on 89 test WSIs with exhaustive annotations of lung cancer metastasis. Qualitative and quantitative results are presented in Figure 3 and Table 2, respectively. In qualitative comparisons, no significant difference was observed between GMP and ReLSE pooling. However, self-training markedly enhanced lesion recall. Quantitative analysis revealed that both ReLSE and self-training significantly enhanced localization performance across ITC, micro, and macro lesions. Integrating ReLSE and self-training with streaming CNN notably improved overall recall (72.27% vs. 42.98%), as well as a recall for ITC (76.32% vs. 26.41%), micro (76.11% vs. 41.68%), and macro lesions (71.62% vs. 43.36%) compared to the baseline model. While our approach still falls short of a strongly supervised FPN model trained on 413 exhaustively annotated slides for overall recall (72.27% vs 81.80%), it achieved markedly higher sensitivity for ITC (76.32% vs. 51.73%) and micro lesions (76.11% vs. 72.74%).

**Table 2:** Lesion Localization Performance for Lung Cancer Metastasis in Lymph Nodes. Four weakly supervised streaming CNN models were trained on 3,024 LN images with LN-level labels. Two semi-supervised FPN mixed pseudo labels from ReLSE-CAM-FPN with 20% and 100% of manual annotations, respectively. A strongly supervised FPN was trained using manual annotations. These models were evaluated on 89 test WSIs with exhaustive annotations. Each patch measures 64 × 64 μm^2^. Patch-level recall is defined as the ratio of true positive patches to the total number of positive patches. A lesion-level recall is determined by the presence of any predicted positive patches within a lesion area, categorizing it as a predicted positive lesion.

|  |  | Patch-level Recall |  |  |  | Lesion-level Recall |  |  |
| --- | --- | --- | --- | --- | --- | --- | --- | --- |
| Model | Precision | All | ITC | Micro | Macro | ITC | Micro | Macro |
| Streaming CNN (Weakly supervised learning) |  |  |  |  |  |  |  |  |
| GMP-CAM (baseline) | 0.8000 | 0.4298 | 0.2641 | 0.4168 | 0.4336 | 0.5179 | 0.9069 | **1.0000** |
| ReLSE-CAM | 0.8000 | 0.5346 | 0.4426 | 0.5328 | 0.5358 | 0.7245 | 0.9441 | **1.0000** |
| GMP-CAM-FPN | 0.8000 | 0.7233 | 0.6260 | 0.7570 | 0.7189 | 0.6529 | 0.9388 | **1.0000** |
| ReLSE-CAM-FPN (proposed) | 0.8000 | 0.7227 | 0.7632 | 0.7611 | 0.7162 | 0.8182 | 0.9574 | **1.0000** |
| Semi-supervised learning |  |  |  |  |  |  |  |  |
| FPN (20% annotations; proposed) | 0.8000 | 0.8151 | 0.7596 | 0.8168 | 0.8154 | 0.8230 | 0.9563 | 0.9859 |
| FPN (100% annotations; proposed) | 0.8000 | **0.9112** | **0.8002** | **0.9033** | **0.9135** | **0.8705** | **0.9681** | 0.9867 |
| Strongly supervised learning |  |  |  |  |  |  |  |  |
| Strongly supervised FPN | 0.8000 | 0.8180 | 0.5173 | 0.7274 | 0.8356 | 0.6143 | 0.8856 | 0.9467 |

**Figure 3:**
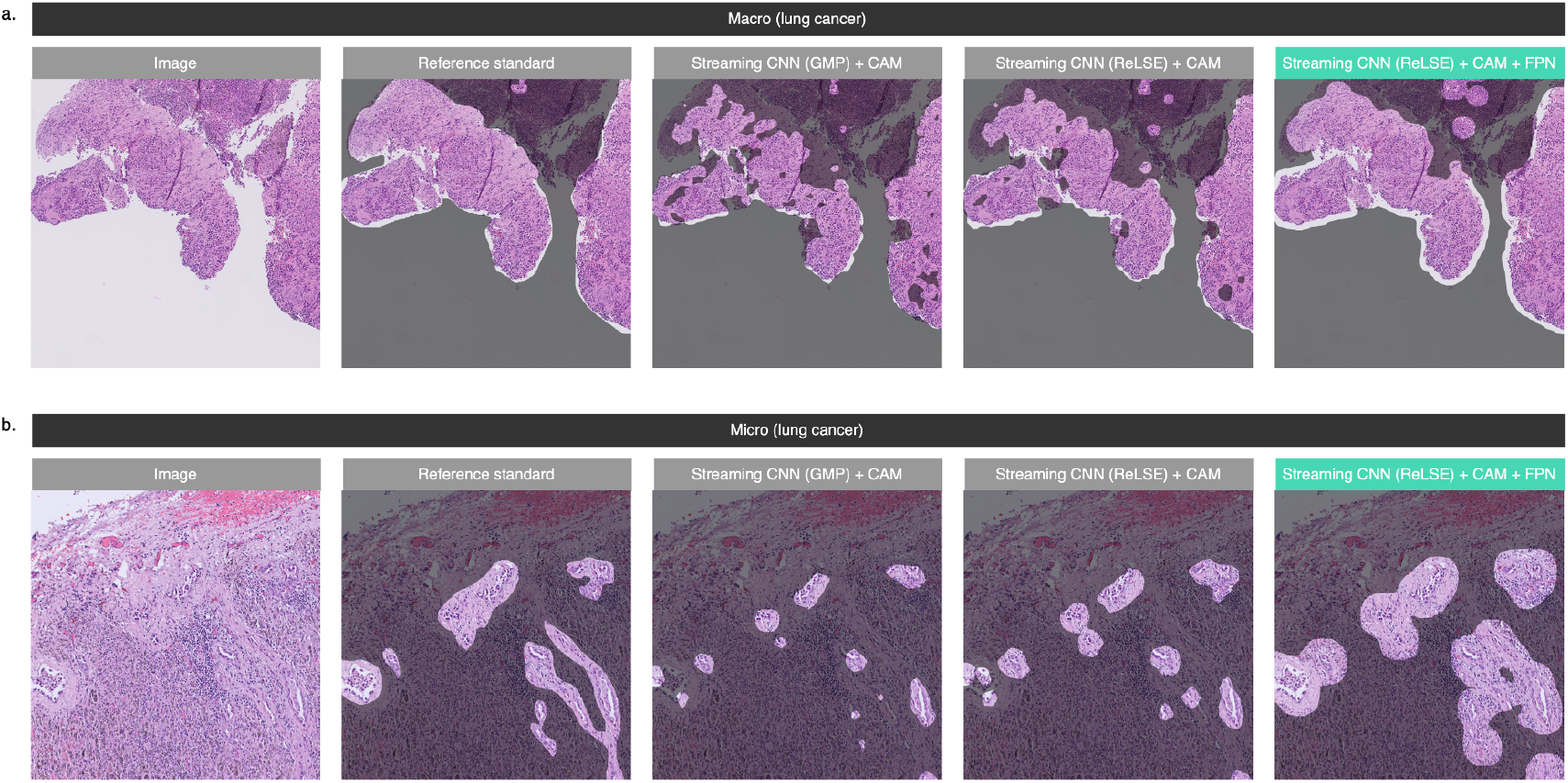
Lung cancer LN metastasis lesion localization of various models on the TVGH dataset. (a) Image of macro and the identification results of various models. (b) Image of ITC.

To further enhance performance, we combined detailed annotations with pseudo labels to train semi-supervised models by merging contours. Using the full set of detailed annotations, the semi-supervised model outperformed the strongly supervised model, achieving a higher overall recall (91.12% vs. 81.80%), as well as improved recall for ITC (80.02% vs. 51.73%), micro (90.33% vs. 72.74%), and macro lesions (91.35% vs. 83.56%).

## Discussion

Recent advancements in artificial intelligence have shown that leveraging large and diverse datasets can significantly enhance model performance, often more effectively than intricate algorithmic designs. This paper introduces a novel method that improves both classification and lesion localization without relying on explicit lesion annotations. This approach not only reduces the associated costs but also allows models to learn from a larger volume of data, facilitating the development of AI-assisted pathology assessment workflows. In daily pathological diagnosis, a wide spectrum of diseases is encountered, ranging from non-neoplastic processes to common cancers and rare malignancies. To ensure that a model captures all potential normal and abnormal features, large datasets are indispensable. However, the necessity of such datasets presents significant challenges for manual annotation. Beyond disease diagnosis and classification, precise lesion localization on pathology slides is equally critical. Many diagnostic tasks, such as tumor size measurement, rely on accurate localization. Additionally, precise lesion localization enhances visual explainability, allowing pathologists to verify the AI model’s inferences or predictions visually. This capability fosters greater confidence in adopting AI applications within diagnostic workflows.

In this study, we propose an efficient solution for developing AI-assisted digital pathology applications. Our experimental results demonstrate that the proposed method achieves state-of-the-art performance in both classification and lesion localization tasks without requiring detailed annotations. Furthermore, our method is flexible enough to incorporate detailed annotations, enabling semi-supervised training. Even with limited detailed annotations, the proposed method achieves performance comparable to that of models trained using strongly supervised learning, highlighting its potential to balance annotation efficiency with diagnostic accuracy.

A current limitation of the streaming convolution algorithm and ReLSE pooling lies in their incompatibility with vision transformers^24–26^. Although vision transformers do not currently exhibit substantial superiority over CNNs in tasks involving regular-sized datasets, they have shown greater capacity for learning from large datasets. Many foundation models for natural images^27,28^ and pathology^29–34^ have adopted vision transformer architectures, demonstrating strong generalization capabilities across various downstream tasks. The ViT^24^, a conventional vision transformer, is inherently incompatible with the streaming algorithm due to its unlimited receptive fields in every self-attention layer. However, Swin-based architectures^25,26^, which have limited receptive fields in their early layers, can be streamly computed to reduce memory footprint. When extending Swin architectures to gigapixel-sized images, the final global average pooling (GAP) layer can be replaced by ReLSE in a drop-in manner, mitigating the issue of gradient vanishing.

Ultimately, the goal is to develop foundation models for pathology that eliminate the need for task-specific training in the future. Most current literature employs a self-supervised learning paradigm to learn image features by distinguishing one image patch from another, relying solely on image data.^29–33^ We hypothesize that integrating vision and text modalities can further advance foundation models for pathology. Text embeddings capture complex relationships, such as the hierarchical connection between malignancy and carcinoma, and the progressive nature of conditions from mild dysplasia to severe dysplasia, carcinoma in situ, and invasive carcinoma. An effective training pipeline should, therefore, learn both similarities and distinctions—not just differences. Moreover, a foundation model aligned with a text encoder could be used in text-related applications, such as reasoning and report generation. To develop vision-language models, our method can be applied as a weakly supervised learning technique for whole-slide images, leveraging WSIs and de-identified medical records, whether in plain text or structured data, which are typically associated with a study or WSI.

## Methods

### Datasets

For the identification of lymph node (LN) metastasis in breast cancer, we utilized the publicly available CAMELYON16 dataset^18^, comprising 111 positive and 159 negative whole-slide images (WSIs) for training, and 48 positive and 81 negative WSIs for testing. To enhance stratified evaluation, we incorporated the CAMELYON17 dataset^18^, which includes 36 ITC, 59 micro, 87 macro, and 318 negative WSIs. For lung cancer classification tasks, we employed the TCGA-LUAD and TCGA-LUSC datasets, consisting of 530 WSIs for adenocarcinoma and 513 for squamous cell carcinoma.

Additionally, we retrospectively analyzed slides from surgical resections of lymph nodes in lung carcinoma dissections, collected between 2020 and 2021 from the pathology archive at Taipei Veterans General Hospital (TVGH). This cohort encompassed 140 cases (116 for training and 24 for test) and 502 hematoxylin and eosin (H&E) slides (413 for training and 89 for test). 3,024 lymph nodes on training slides were annotated. The WSIs were scanned and digitized at 40× magnification (0.25 µm/pixel) using an IntelliSite Ultra Fast Scanner (Philips, Amsterdam, Netherlands). The study protocol was reviewed and approved by the Institutional Review Board (IRB) of TVGH (IRB No. 2020-05-006AC), and the requirement for written informed consent was waived by the IRB due to the use of de-identified digital slides.

### ReLSE pooling

In the context of WSIs, identifying small lesions amidst billions of pixels presents a significant challenge. While GAP is widely adopted in CNNs for natural image processing, it has been shown to be inadequate for WSIs. GAP tends to dilute lesion features by incorporating information from irrelevant regions, leading to the gradient vanishing issue, which undermines effective learning.^17^ In contrast, GMP addresses the issue of sparsity by focusing on the most representative regions of WSIs, making it a common choice for WSI processing in various approaches, such as conventional MIL, MIL variants employing feature extractor pretraining (e.g., MIL-RNN^8^), whole-slide training methods^17^, and streaming CNNs^1,15^.

However, lesions in WSIs often span multiple regions, and GMP restricts gradient flow to only the most prominent regions, thereby limiting the utilization of the entire lesion information. This inefficient use of data hinders the model’s ability to fully capture the complexity of the lesion’s spatial distribution. To overcome both the gradient vanishing problem and the inefficiency in data utilization, we introduce Rectified LogSumExp (ReLSE) pooling, a method designed to enhance feature learning by balancing the focus on both representative and sub-representative regions, thereby improving model performance in WSI-based tasks.

Specifically, let *Z* ∈ (ℛ^+^)^*H*×*W*×*C*^ represent the embedding feature map produced by the convolutional layers, where *H, W*, and *C* denote the height, width, and number of channels, respectively. We use *z*_*ijc*_ ∈ ℛ^+^ to indicate the value in *Z* at position (*i, j, c*), where *i* ∈ [0, *H*), *j* ∈ [0, *W*), and *c* ∈ [0, *C*). The conventional GAP and GMP can be expressed as,

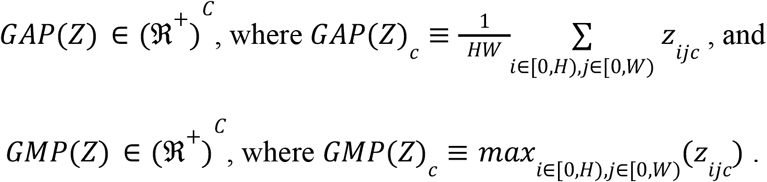

Our proposed ReLSE is formulated as:

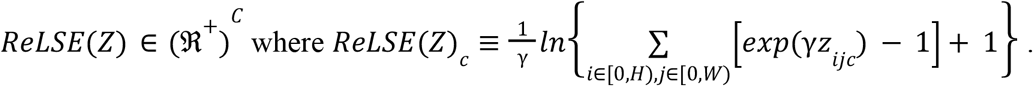

Here, γ ∈ ℛ^+^ is a hyperparameter controlling the smoothness of the operation. Figure 4 illustrates the relationship among ReLSE, GMP, and GSP. In all experiments, we set γ = 1 to balance smoothness and feature retention.

**Figure 4:**
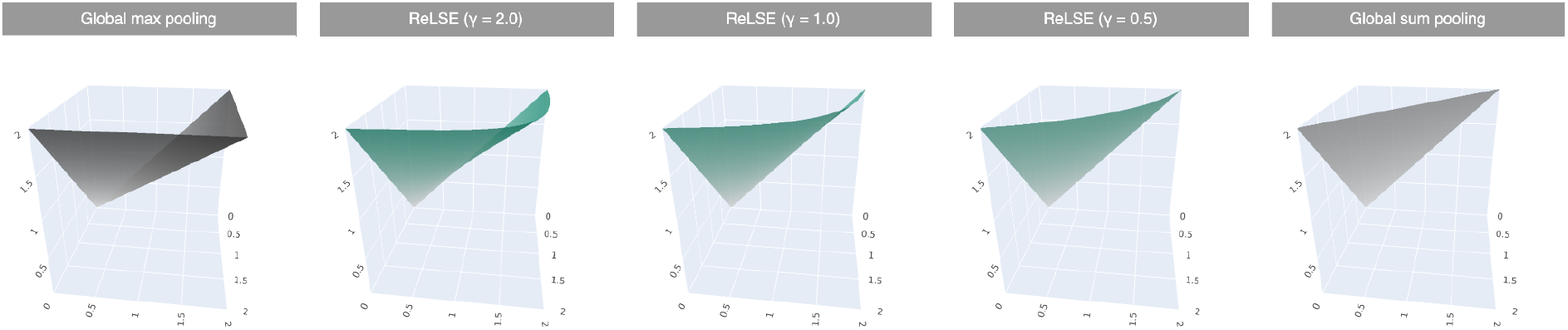
Illustration of pooling operations. Given two independent variables, x and y, as values in embedding feature maps, the outputs of different pooling operations are plotted as z.

### Properties of ReLSE pooling

The ReLSE function establishes a lower bound with respect to the GMP function. Let *z*^∗^= *GMP*(*Z*)_*c*_ represent the maximum, or one of the maxima in the case of multiple maximum values, among *z*_*ijc*_. This relationship can be expressed as follows:

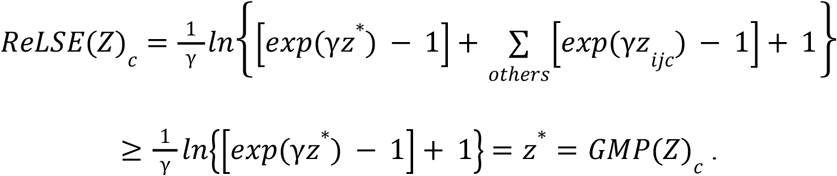

Since ∀*z* ≥ 0, [*exp*(*z*) − 1] ≥ 0 and *ln*(·) is an increasing function, the inequality holds.

Additionally, the ReLSE function is bounded from above by global sum pooling (GSP). In natural image classification, where images are resized to a fixed dimension before model input, GSP is functionally equivalent to global average pooling (GAP) aside from a scale factor, which can be compensated by adjusting the weights in the final linear layer. However, in WSI classification, GSP and GAP differ due to the variable size of WSIs. We hypothesize that GSP is more suitable than GAP, based on the assumption that larger images encapsulate more information, thereby making the model more confident in its predictions, as opposed to merely diluting the existing information. GSP amplifies the embedding values as the image size increases, pushing the predicted probabilities closer to either 0 or 1. Accordingly, we define the ReLSE function as being bounded by GSP, demonstrated by the following inequality:

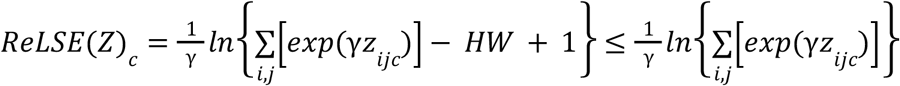

since (− *HW* + 1) ≤ 0 and *ln*(·) is an increasing function. By leveraging the convexity of *exp*(·), we derived a second inequation:

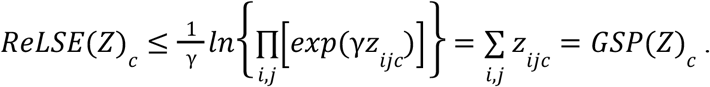

The smoothness hyperparameter γ can be tuned to shift ReLSE toward either GMP or GSP. When γ → ∞, ReLSE converges to GMP. Let *z*^∗^= *GMP*(*Z*)_*c*_, as shown by applying L’Hôpital’s rule,

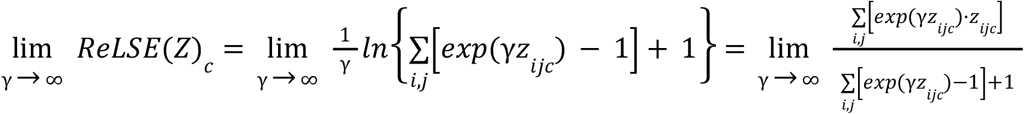

if the right-hand side (RHS) converges. The RHS fraction can be simplified by reducing 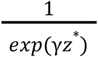:

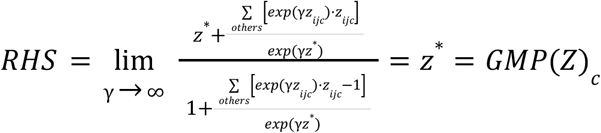

which converges. Thus, the following formula holds:

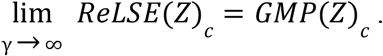

Conversely, when γ → 0^+^, ReLSE converges to GSP. This behavior is similarly demonstrated using L’Hôpital’s rule:

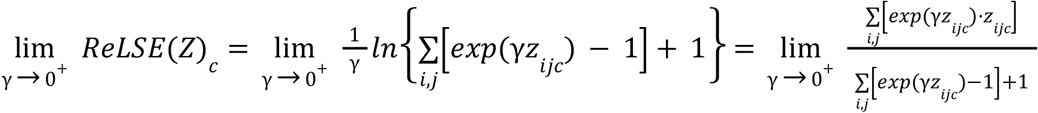

if the RHS converges. The RHS can be simplified as,

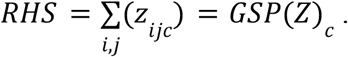

Therefore, ReLSE converges to GSP when γ → 0^+^.

### Self-training with adaptive pseudo annotation generation

Class Activation Mapping (CAM) is a widely adopted technique for segmenting relevant regions in a classification model under weak supervision. However, it tends to emphasize only the most representative features, often neglecting the broader context. This approach is inherent to classification models as they are optimized to distinguish between signal and noise, including potentially informative but less reliable features. To address this limitation, self-training has proven effective. In this approach, CAM or its variants, such as GradCAM^35^, are utilized as pseudo-annotation generators to train a semantic segmentation model. This segmentation model then aims to enhance pixel-wise accuracy by aligning its optimization objectives accordingly.

In practice, assigning pseudo annotations requires setting an appropriate threshold to differentiate between positive and negative regions on a CAM map. This task becomes particularly challenging in streaming CNNs, where the distribution of CAM activation values can vary significantly across tasks, class distributions, and pooling layers.

To address this challenge, we propose a method for adaptively determining the threshold. We observed distinct distributions of CAM activation values corresponding to lesion and non-lesion areas. The activation values associated with lesions were spread across a wide range, with some even overlapping with the distribution of non-lesion areas. This indicates that CAM may not uniformly highlight lesion areas. In contrast, the activation values for non-lesion areas were more constrained within a limited range. Leveraging the distribution of CAM activation values for non-lesion areas thus provides a robust means of distinguishing between lesion and non-lesion regions.

Our algorithm first fits a Gaussian model to the CAM activation values associated with negative tissues in the validation dataset. The threshold is then determined using the formula μ + kσ, where μ and σ represent the mean and standard deviation of the fitted model, respectively, and k is a hyperparameter that balances sensitivity and specificity. For all our experiments, we set k to 3. Subsequently, CAM maps are inferred from the training images, which are then employed to generate pseudo annotations. Finally, a segmentation model, such as the Feature Pyramid Network (FPN) used in our experiments, is trained using pairs of training images and their corresponding pseudo annotations. An example is presented as Figure 5.

**Figure 5:**
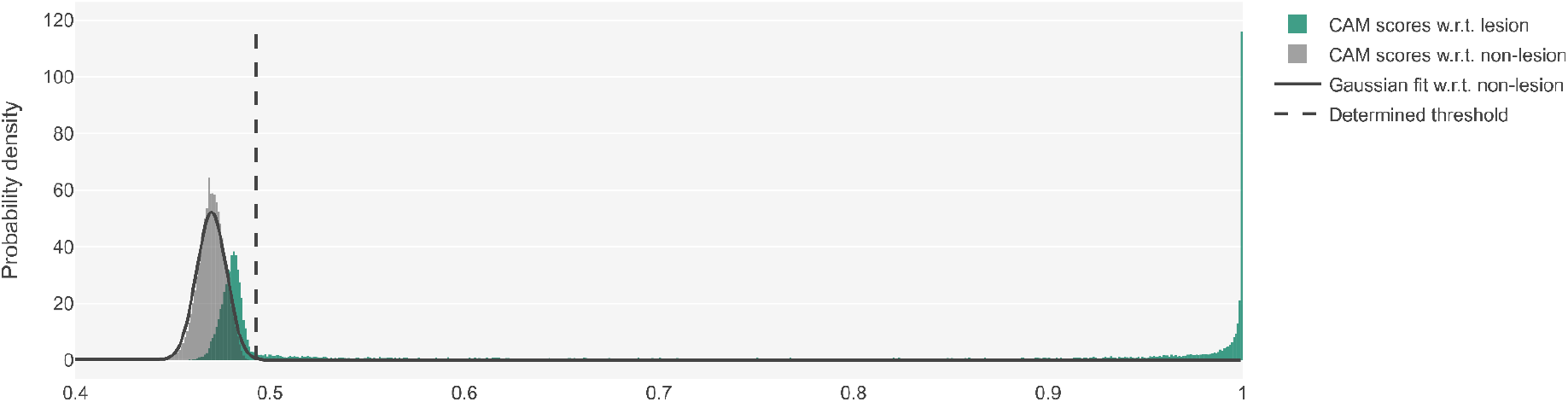
Distributions of CAM scores with regard to lesion and non-lesion areas and our proposed adaptive thresholding algorithm. These data were generated by a streaming CNN trained with ReLSE pooling and the CAMELYON16 dataset. The Gaussian model was fitted using CAM scores in negative validation WSIs. The threshold was determined by calculating μ + 3σ.

### Experimental setups and statistics

This section outlines the experimental setup in detail. Model training and inference were executed using 8 NVIDIA GeForce RTX 3090 GPUs running in parallel. Both the streaming CNNs and FPNs were initialized with pre-trained weights from the ImageNet Large Scale Visual Recognition Challenge (ILSVRC) 2012 dataset (ImageNet-1k)^28^. During training, each GPU processed one image per step. For the breast cancer lymph node metastasis task and lung cancer type classification, images at a resolution of 0.25 μm/pixel were downscaled to 0.5 μm/pixel, while for the lung cancer lymph node metastasis task, images were downscaled to 1.0 μm/pixel. To augment the training data, we applied random flipping, random rotations (−180° to 180°), stain matrix perturbations^36^ (−10° to 10°), and variations in stain concentrations (0.33× to 3.0×) before the images were fed into the training pipeline. The streaming CNNs were optimized using the AdamW optimizer^37^ with an initial learning rate of 1e−5, beta values of [0.9, 0.999], and a weight decay of 0.01 to minimize the cross-entropy loss. Validation losses were monitored after each epoch, and the model with the lowest validation loss was saved if no improvement was observed for four consecutive epochs. FPNs were updated using stochastic gradient descent with an initial learning rate of 0.01, a momentum of 0.9, and L2 decay of 0.0005, following a polynomial decay learning rate schedule (power: 0.9, iterations: 80,000). The software stack used for this process included CUDA 11.8 for GPU acceleration, PyTorch 2.0.1, mmsegmentation 1.1.2, and Torchvision 0.15.2 for model construction and training, Open MPI 4.1.2 and Horovod 0.28.1 for multi-GPU training, OpenCV 4.10.0.84 and Pillow 9.5.0 for image processing, scikit-learn 1.5.1 for Gaussian model fitting, and OpenSlide 3.4.1 for WSI decoding. The confidence intervals (CIs) of AUCs were calculated using the Delong method (pROC 1.18.0).

## Acknowledgments

The results here are in whole or part based upon data generated by the TCGA Research Network: https://www.cancer.gov/tcga.

## Supplementary

### Classification of lung cancer types

We utilized the TCGA-LUAD and TCGA-LUSC datasets, comprising 530 lung adenocarcinoma specimens and 513 squamous cell carcinoma specimens, to evaluate the classification performance of our method against other weak-supervised alternatives. The dataset was randomly partitioned into 667 slides for training, 166 for validation, and 210 for testing. The baseline streaming CNN with GMP achieved an AUC of 0.9789 (95% CI: 0.9631–0.9948). In comparison, the streaming CNN with ReLSE pooling reached an AUC of 0.9750 (95% CI: 0.9592–0.9908). While the performance difference between the two models was not statistically significant, both demonstrated competitive discriminative ability compared to existing approaches in other literature, including DTFD-MIL (AUC: 0.961, MaxMinS, ResNet-50, 0.5 μm/pixel)^19^ and IBMIL (AUC: 0.9254, Trans-MIL, ResNet-18, 0.5 μm/pixel)^20^. Due to the lack of ground truth detailed annotations in the test set, quantitative analysis of lesion localization abilities cannot be conducted.

